# Medial septum glutamatergic neurons drive speed coding and grid cell spatial accuracy in the medial entorhinal cortex

**DOI:** 10.64898/2026.09.28.754654

**Authors:** Jennifer Robinson, J. Eric Carmichael, Mark P. Brandon, Michael E. Hasselmo

## Abstract

Successful navigation requires the integration of self-motion signals with spatial representations, yet the circuits that provide velocity information necessary for stable grid cell firing remain incompletely understood. Here, we identify medial septal (MS) glutamatergic neurons as a critical component of this pathway. Using cell-type-specific optogenetic silencing combined with in vivo electrophysiology in freely moving mice, we show that MS glutamatergic neurons support the fidelity of firing rate–based speed coding in the medial entorhinal cortex (MEC). Silencing this population reduced grid cell spatial periodicity and stability and produced distortions of grid firing fields. Silencing also reduced theta phase-locking strength in strongly theta-modulated MEC neurons. Model simulations of a hybrid oscillatory interference–continuous attractor network with reductions in speed-signal fidelity disrupted grid periodicity and reproduced features of the spatial distortions observed in vivo. Together, these findings identify a septo-entorhinal circuit that coordinates self-motion information with the spatial and temporal organization of MEC activity to maintain a stable neural representation of space.

**HIGHLIGHTS:**

- Medial septum glutamatergic neurons support the fidelity of speed coding in the medial entorhinal cortex
- Silencing MS glutamatergic neurons distorts grid periodicity and spatial stability
- Medial septum glutamatergic input regulates MEC theta phase locking
- Model simulations identify speed-signal fidelity as important for grid periodicity

## INTRODUCTION

Successful navigation requires the brain to construct an internal representation of space, referred to as a cognitive map. A central component of this map is the medial entorhinal cortex (MEC), which contains many distinct cell types that code for various aspects of space^1–4^. This includes grid cells which exhibit striking hexagonal firing patterns that tile the environment and are predicted to provide a regular metric for spatial navigation and memory^1,5–9,10^.

Multiple computational frameworks including continuous attractor network (CAN), oscillatory interference (OI), and hybrid models have been proposed to explain grid cell firing ^8,11,12,13^. Despite their differences, these models converge on a central requirement: the integration of velocity signals. In CAN models, velocity inputs drive shifts in the activity bump through conjunctive grid, speed, and head-direction signals to perform translational path integration. In OI models, the burst frequency of velocity-controlled oscillators (VCOs) has a linear dependence on running speed. Consistent with these predictions, the MEC contains several cell types that display changes in their activity based on their running speed^2,14–18^, including principal cells that exhibit increased firing rates as a function of running speed^15,1617^. Disruption of self-motion signals, such as during passive transport, abolishes grid cell periodicity, reinforcing the importance of velocity inputs for spatial coding^19^.

The speed signals in the MEC arise from multiple circuits. Speed information is conveyed through an ascending pathway from the pedunculopontine nucleus through the horizontal limb of the diagonal band to the MEC^20^, while local MEC circuitry further shapes speed coding, with parvalbumin-expressing interneurons contributing to both speed modulation and grid cell spatial periodicity^20,21^. Other subcortical structures may provide additional locomotor signals including the supramammillary nucleus, where neurons encode future locomotor speed and selectively regulate speed-modulated hippocampal activity^22^.

The activity of the MS is closely linked to velocity coding. The MS acts as a central node for both rhythmic and velocity-related input to the hippocampal-entorhinal network. Septal neurons exhibit robust speed modulation, expressed through both firing rate increases and modulation of theta-rhythmic bursting as a function of running speed^23–25^. It is well established that the frequency of theta oscillations throughout the hippocampal-entorhinal network are positively modulated by running speed^25,14,26^. Notably, two dissociable running speed signals have been identified in MEC; firing rate speed coding and intrinsic theta frequency speed coding^17,27^. Pharmacological inactivation of the MS disrupts grid cell spatial periodicity^27^ and also perturbs speed-related signals in the MEC^17^ supporting a role for septal inputs in velocity-dependent coding.

The MS is composed of diverse neuronal populations, including cholinergic, GABAergic, and glutamatergic neurons that differ in their molecular identity, connectivity, and physiological properties^28^. Chemogenetic modulation of cholinergic activity has revealed that MS cholinergic neurons have a moderate influence on theta dynamics without disrupting grid periodicity^29^. MS GABAergic neurons that project to the entorhinal cortex display rhythmic firing during locomotion events^30,31^, consistent with a role in coordinating network oscillation. While optogenetic pacing of PV neurons does not disrupt grid cell spatial firing, even at frequencies well above the theta range (30 Hz)^32–35^, while optogenetic inhibition of GABAergic septal populations reduces entorhinal theta power and disrupting hexagonal grid cell periodicity^36^, indicating a critical role in maintaining spatial coding.

MS glutamatergic neurons are a heterogeneous population consisting of distinct activity and projection patterns to different regions^37,38^ and project to many cortical and subcortical regions^39,40^, including long-range excitation to both interneurons and pyramidal cells in the MEC^41,42^. Consistent with this heterogeneity, septal glutamatergic neurons have been linked to several different functions, including rhythm generation, locomotion initiation, and whisking activity^39,40,43^. Rhythmic optogenetic activation of this population can robustly drive theta oscillations^39,40^, yet optogenetic silencing produces no significant changes in theta power or frequency^44^, suggesting a more nuanced role in network dynamics. Calcium imaging during exploration identifies three distinct activity clusters within MS glutamatergic neurons, with approximately one-third selectively active during locomotion^38^. In addition, MS glutamatergic terminal activity in the MEC has been linked to movement initiation in head-fixed animals^41^, suggesting a potential role in conveying movement-related signals relevant for velocity-dependent coding.

While previous work has characterized septal contributions to theta rhythms and grid cell spatial coding, a key question remains: which MS cell type contributes to the velocity signals required for grid cell spatial firing? In particular, do MS glutamatergic neurons influence speed signals in the MEC, and are these contributions necessary to maintain grid cell spatial periodicity? Here, we address this question by cell-type-specific optogenetic inhibition of MS glutamatergic neurons combined with unit recordings in the MEC. Our results reveal that manipulation of this population alters speed coding and distorts grid cell firing fields, providing new insight into the circuit mechanisms that support spatial representations for navigation.

## Results

### Selective silencing of MS glutamatergic neurons does not alter open-field exploration or MEC theta power

The MS glutamatergic neurons are heavily implicated in theta rhythmicity and speed signaling in the hippocampal-entorhinal network. Therefore, to unpack the contribution of MS glutamatergic neurons in theta rhythmicity and speed coding, we used selective optogenetic silencing together with in vivo recordings in the MEC (Figure 1). To target MS neurons, Archaerhodopsin-3 variant ArchT was expressed in the MS. ArchT was expressed selectively in MS glutamatergic neurons using a Cre-dependent AAV in VGLUT2-Cre mice (Fig. 1Ai). We further examined the connection between glutamatergic neurons and the MEC using retrograde-Arch injected directly into the MEC. While MS glutamatergic terminals have previously been identified in the MEC^39,41,42^, we examined the distribution of MEC projecting cells in the MS and found a distinctive distribution of cell bodies mostly located along the midline in the ventral portion of the MS (Figure 1Aii). We also examined the pattern of MS glutamatergic terminals in the MEC, and observed MS glutamatergic terminal expression across layers of the MEC, with a higher density of projections in superficial layers II/III as well as layer V (Figure 1B).

**Figure 1.**
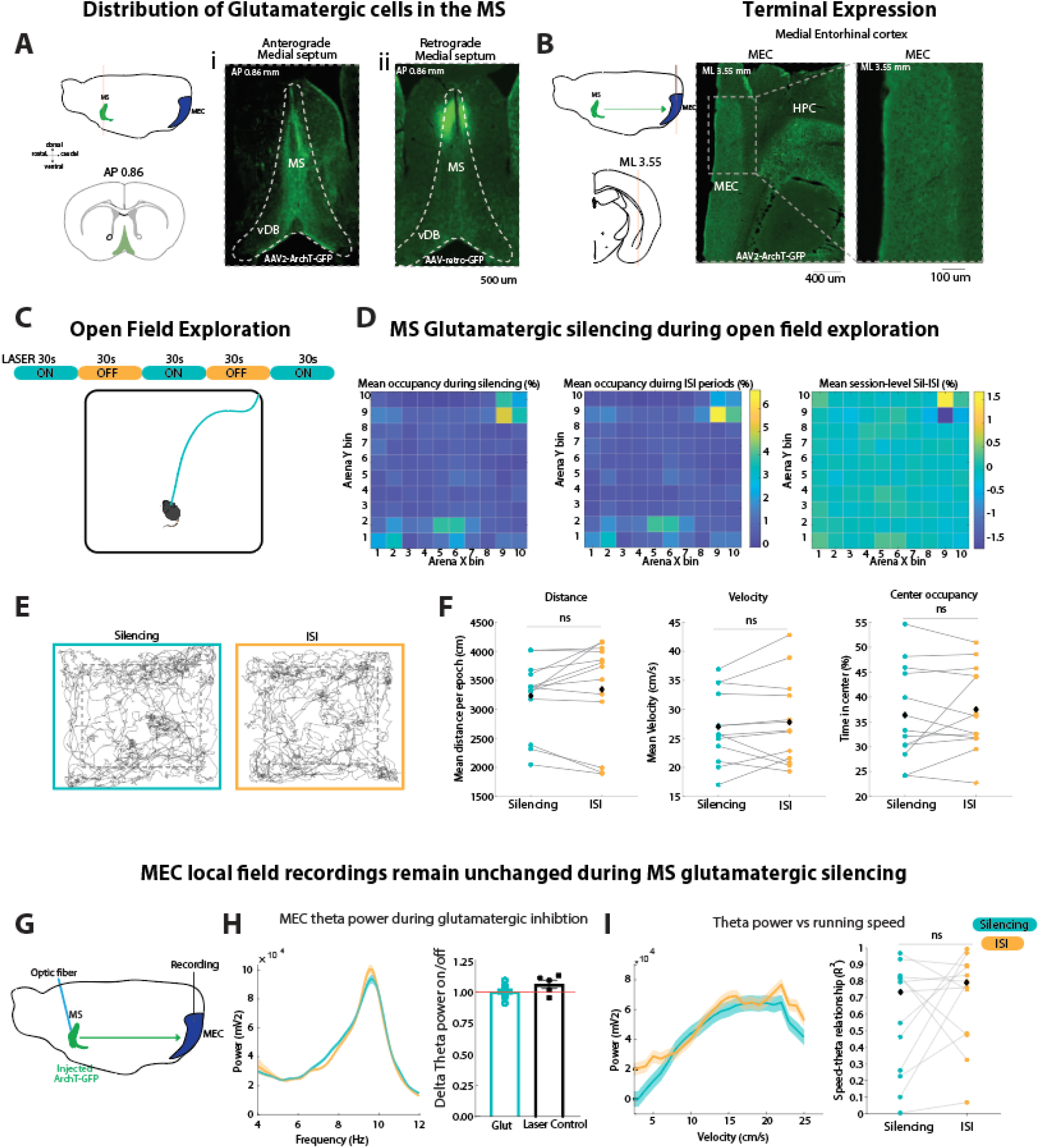
Selective inhibition of medial septal glutamatergic neurons does not alter open-field exploration or MEC local field potential activity. **(A)** Anatomical distribution of medial septal (MS) glutamatergic neurons. Left, schematics showing the locations of anterograde labeling from the MS to the medial entorhinal cortex (MEC; top) and the corresponding coronal level of the MS (bottom). Right, (i) representative coronal sections showing anterograde labeling following injection of AAVdj-ArchT-GFP into the MS and (ii) retrograde labeling of MS glutamatergic neurons projecting to the MEC. Dashed lines delineate the MS. Scale bar, 500 μm. **(B)** Terminal expression of MS glutamatergic projections in the MEC. Left, schematic showing the MS-to-MEC projection and approximate coronal level examined. Middle and right, representative images showing ArchT-GFP-positive axonal terminals in the MEC and a higher-magnification view of terminal labeling, respectively. Dashed lines indicate anatomical boundaries. Scale bars, 400 μm (middle) and 100 μm (right). **(C)** Open-field exploration paradigm. Mice freely explored a square arena during alternating 30-s periods of optogenetic silencing (laser ON; cyan) and inter-silencing interval (laser OFF; orange). **(D)** Spatial occupancy during MS glutamatergic silencing. Heatmaps show mean occupancy across recording sessions during silencing (left) and inter-silencing (ISI) (middle) periods. Right, session-level difference in spatial occupancy between silencing and ISI periods (silencing − ISI). **(E)** Representative locomotor trajectories during MS glutamatergic silencing (cyan) and ISI (orange) periods. **(F)** Mean distance traveled per epoch (left), mean running velocity (middle), and percentage of time spent in the center of the arena (right) during MS glutamatergic silencing and ISI periods. Connected points indicate paired measurements from individual recording sessions; black diamonds indicate group means. ns, not significant. **(G)** Experimental configuration for simultaneous optogenetic inhibition of MS glutamatergic neurons and local field potential (LFP) recordings in the MEC. AAV-ArchT-GFP was injected into the MS, an optic fiber was positioned above the MS, and LFPs were recorded from the MEC. **(H)** MEC theta oscillations during MS glutamatergic silencing. Left, mean LFP power spectra during silencing (cyan) and ISI (orange) periods. Shading indicates SEM. Right, theta power during MS glutamatergic silencing relative to ISI periods for MS glutamatergic (Glut) and laser-control recordings. Individual points indicate recording sessions; bars show mean ± SEM. **(I)** Relationship between running speed and MEC theta power during MS glutamatergic silencing. Left, mean theta power as a function of running velocity during silencing (cyan) and ISI (orange) periods; shading indicates SEM. Right, coefficient of determination (R²) for the relationship between running speed and theta power during silencing and ISI periods. ns, not significant.

The stimulation protocol used throughout the recording session consisted of a 30 second laser on periods to silence glutamatergic activity followed by 30 second inter-stimulus interval (ISI) periods repeated throughout the entire recording session (Figure 1C). Given previous reports that have indicated that silencing MS glutamatergic neurons reduces animals’ exploration of open arms of an elevated plus maze^43^, we examined if silencing has an effect on the animals’ exploration of a familiar open field environment (Figure 1D-F). We first examined mean occupancy of the open field during MS glutamatergic silencing versus ISI periods by normalizing the animals’ occupancy to the entire recording session and found no pattern of exploration aside from a preference for one quadrant of the open field, which was consistent across both the silencing and ISI periods and was not significantly different from normalized bins (*p* > 0.05; Figure 1D). Next, we compared silencing with ISI periods and found no change in distance traveled (Sil, 3216 ± 184.4 cm; ISI, 3291 ± 234.9 cm; paired *t* test, *p* = 0.4745), mean velocity during exploration (Sil, 27.11 ± 6.133 cm/s; ISI, 28.02 ± 7.071 cm/s), or occupancy in the central 50% of the open field (Sil, 10.68 ± 2.956 min; ISI, 10.95 ± 2.481 min; two-tailed paired *t* test, *p* = 0.5534; Figure 1F). Wilcoxon signed-rank tests for distance, speed, and center occupancy yielded *p* = 0.414, *p* = 0.455, and *p* = 1.00, respectively.

The MS is the central driver of hippocampal-entorhinal theta oscillations; however, recent evidence suggests that while optogenetic activation of MS glutamatergic neurons can drive theta activity, optogenetic silencing of MS glutamatergic neurons has no effect on either theta power or frequency^44^. Consistent with current literature, LFP recordings from the superficial layers of the MEC showed no change in theta-band activity during MS glutamatergic silencing relative to ISI periods (Glut, 1.005 ± 0.0144; Control, 1.0630 ± 0.3300, mean ± SEM; Figure 1H). It is well established that theta oscillations throughout the hippocampal-entorhinal network are positively modulated by running speed^26,45,46^. Previous reports have shown that optogenetic activation of MS glutamatergic neurons drives both theta oscillations and locomotor behavior in head-fixed animals^41^. During open-field exploration, silencing MS glutamatergic neurons did not significantly alter the strength of the relationship between running speed and theta power (Sil, R² = 0.1998 ± 0.2011; ISI, R² = 0.4863 ± 0.1521; paired Wilcoxon signed-rank test, *p* = 0.2440; Figure 1i).

To confirm that each recording session that the applied light was sufficient to drive an optogenetic response, we performed 100-150 square pulse stimulation of 15 ms at 1 Hz following every experimental recording and calculated the stimulation triggered average on the LFP (See supplemental figure 1E,I). Interestingly, MS glutamatergic neurons result in the appearance of a bimodal or polysynaptic response, with first a peak at 0.5 sec following the stimulation followed by a trough around 0.6 sec. These results could be due to a mix of direct projections from the MS Glutamatergic neurons directly to the MEC as well as indirect targeting through intraseptal connections to adjacent MS GABAergic neurons^40^. Together, these results indicate that silencing MS glutamatergic neurons does not detectably alter MEC theta power, and does not cause changes in locomotor activity or trajectories compared to controls when mice are free foraging in an open field environment.

### MS glutamatergic neurons support speed coding in the MEC

It has been well characterized that there are multiple speed signals in the MEC^16–18^. The MS supports speed signals to the MEC, as the absence of MS input results in an enhancement of firing rate speed coding and a reduction in oscillatory speed coding^17^. To unpack how MS glutamatergic neurons contribute to these speed coding in the MEC, we recorded single units in the MEC as mice foraged for water droplets in an open field arena. The experimental protocol consists of a baseline recording followed by a stimulation recording session using the repeated 30s silencing periods followed by 30s ISI periods. To assess if MS glutamatergic neurons contribute to speed coding of individual units in the MEC, we examined the firing rate to running speed relationship across all three conditions (Figure 2A-C). Speed cell inclusion criteria used the Kropff et al., 2015 speed score method to select cells that are modulated by the animals running speed (see methods). Silencing MS glutamatergic neurons altered the spatial firing patterns and reduced the fidelity of MEC speed coding activity and running speed (Figure 2AC). Of 44 cells recorded, 22 cells passed criteria for speed cells in the baseline condition. In line with result from Dannenberg et al., 2019, silencing did not cause a significant change to the speed correlation (Mean Baseline 0.0518 ± 0.0106, Silencing 0.0466 ± 0.013, ISI 0.0463 ± 0.0116, Friedman test with Dunn’s multiple comparisons *p =* 0.8338 Figure 2D) nor to the slope of the relationship between firing rate and running speed (Mean Baseline 0.03255 ± 0.0092, Silencing 0.0294 ± 0.0079, ISI 0.0285 ± 0.0073, Friedman test with Dunn’s multiple comparisons *p = 0.5796* Figure 2E), however, we did observe a reduced firing rate vs running speed model-R² (Mean Baseline 0.5454 ± 0.060, Silencing 0.4087 ± 0.0631, ISI 0.4303 ± 0.0079, *Friedman* test with Dunn’s multiple comparisons *p* < 0.01, Base-Sil *p* < 0.01, Base-ISI *p* < 0.05 Figure 2G). The reduced speed model R² suggests an overall reduced precision in speed coding across the population. Together, these results indicate that MS glutamatergic activity supports the fidelity of MEC speed coding.

**Figure 2:**
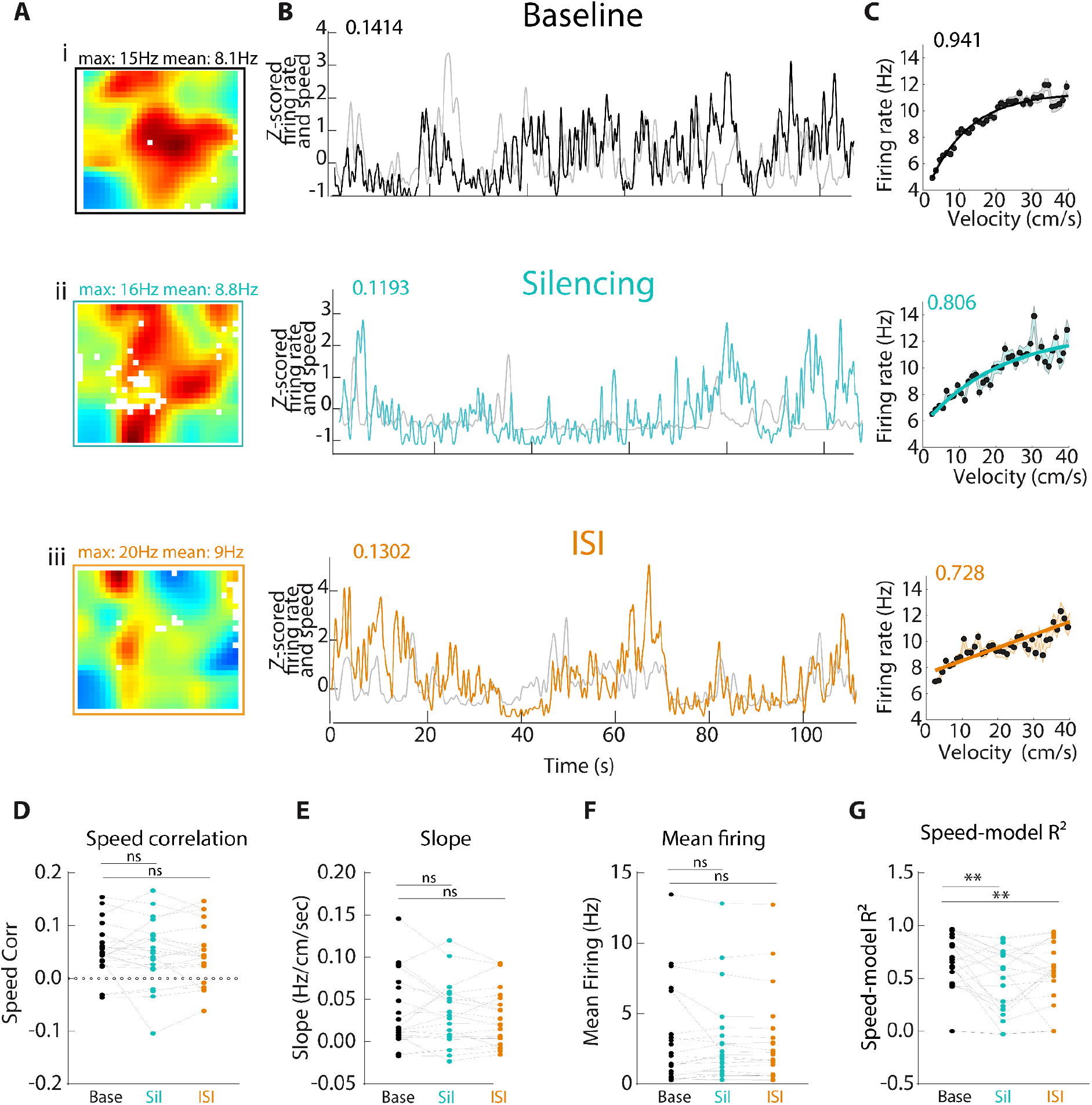
MS glutamatergic neurons support speed coding in the MEC. **(A)** Spatial firing-rate maps of a representative MEC neuron during baseline (i), optogenetic silencing of MS glutamatergic neurons (ii), and the ISI period (iii). Maximum and mean firing rates are indicated above each map. **(B)** Z-scored instantaneous firing rate (black, baseline; cyan, silencing; orange, ISI) and running speed (gray) for the same representative neuron across the recording session. Values above each plot indicate the corresponding speed correlation. **(C)** Firing rate as a function of running velocity during baseline, silencing, and ISI periods for the same neuron. Points indicate firing rates within individual speed bins, and solid lines indicate saturating-model fits. Values above each plot indicate the corresponding saturating-model *R*². **(D)** Number of MEC neurons classified as speed modulated during baseline, silencing, and ISI periods. Colored portions indicate speed-modulated cells and gray portions indicate non-speed-modulated cells; numbers above bars indicate the number of speed-modulated cells. **(E)** Mean firing rate–speed slope during baseline, silencing, and ISI periods. **(F)** Mean firing rate during baseline, silencing, and ISI periods. **(G)** Speed-model *R*² during baseline, silencing, and ISI periods.Values above bars indicate the percentage of cells in each category. Bars in (E) and (F) represent mean ± SEM. **ns**, not significant; **\*\****p* < 0.01; **\*\*\****p* < 0.001.

### MS glutamatergic silencing distorts grid cell spatial coding

Given the importance of speed coding for grid cell spatial coding, we next examine if silencing MS glutamatergic neurons altered grid cell spatial periodicity. To examine this, ratemaps of all silencing periods were concatenated together to visualize spatial dynamics and compared to baseline, the same concatenation was done for ISI periods. We observed that silencing MS glutamatergic neurons results in distinct changes to grid cell spatial periodicity (Figure 3A). Silencing glutamatergic neurons distorts grid cell firing fields during the silencing period, however, gridness during the ISI period was not significantly different from baseline (Baseline gridness, 0.84 ± 0.0696; Sil, 0.3133 ± 0.1125; ISI, 0.5765 ± 0.0849; Friedman test with Dunn’s multiple comparisons, *p* < 0.01; Baseline vs. Sil, *p* < 0.01; Baseline vs. ISI, *p* = 0.1237; Sil vs. ISI, *p* = 0.6620; Figure 3C). Given speed coding is perturbed during glutamatergic silencing, we reasoned that this mismatch in speed signalling may create instability in grid pattern compared to baseline (Figure 3B). Consistent with this, we examined rate map stability by testing the 2D displacement of spatial cross-correlations, which revealed that the overall grid pattern during MS glutamatergic perturbations exhibit reduced spatial stability during both the silencing period and persisted during the ISI periods compared to baseline-sham controls (Baseline-Sil, 3.237 ± 0.7419; Baseline-ISI, 2.320 ± 0.4700; ISI, 1.490 ± 0.2023; Friedman test with Dunn’s multiple comparisons, *p* = 0.01; Baseline-Sil vs. Base-Sham, *p* < 0.0001; Base-ISI vs. Sham, *p* < 0.0001; Sil-ISI vs. Sham,*p* < 0.01; Figure 3D). The reduced stability of the grid cell pattern during silencing and ISI periods was not due to the concatenation of the ratemaps, as shown by sham stimulations which use an equal number of stimulation periods as in the silencing period (Sham controls) as well as controls (Figure S2).

**Figure 3.**
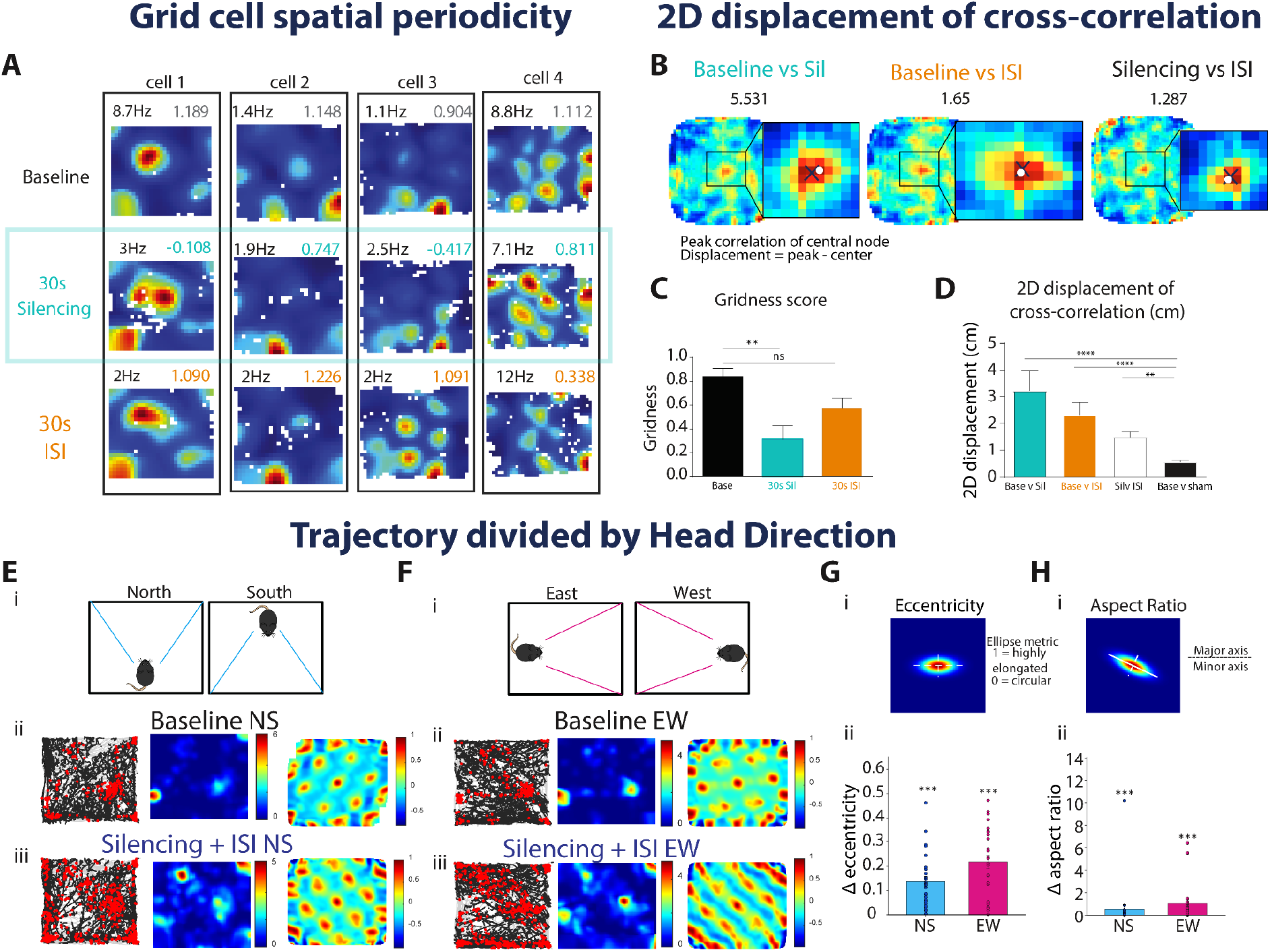
Silencing MS glutamatergic neurons distorts grid cell spatial coding. **(A)** Representative spatial firing-rate maps from four grid cells during baseline, silencing, and the subsequent ISI period. Values above each map indicate mean firing rate (Hz; black) and gridness score (colored). **(B)** Representative two-dimensional cross-correlation maps illustrating displacement of the central correlation peak between baseline and silencing, baseline and ISI, and silencing and ISI periods. Insets show an expanded view of the central autocorrelation peak (x is peak, dot is centroid). Displacement was calculated as the Euclidean distance between the peak correlation and the center of the cross-correlation map. Values above each example indicate peak displacement. **(C)** Mean gridness score during baseline, silencing, and ISI periods. **(D)** Quantification of the two-dimensional displacement of the central cross-correlation peak across experimental comparisons. Bars show group means with error bars; horizontal brackets indicate statistical comparisons. **(E)** Direction-conditioned analysis of grid cell spatial representations for north-and south-directed trajectories. (i) Schematic illustrating the division of trajectories according to northward and southward head direction. (ii–iii) Representative trajectory maps, spatial firing-rate maps, and corresponding spatial autocorrelations during baseline (ii) and silencing (iii). **(F)** Direction-conditioned analysis for east-and west-directed trajectories. (i) Schematic illustrating the division of trajectories according to eastward and westward head direction. (ii–iii) Representative occupancy/trajectory maps, spatial firing-rate maps, and corresponding spatial autocorrelations during baseline (ii) and silencing (iii). Silencing produces a directional elongation of the central autocorrelation structure. **(G)** Quantification of central-peak eccentricity as a measure of autocorrelation-peak elongation. (i) Schematic illustrating eccentricity (for east-west in this schematic example), where values approaching 0 represent a circular peak and values approaching 1 represent an increasingly elongated ellipse. (ii) Magnitude of the change in eccentricity from baseline for north-south (NS) and east-west (EW) directional representations. For each cell, change from baseline was calculated as (Silencing+ISI)−(Baseline), such that 0 represents no change from baseline. **(I)** Quantification of central-peak aspect ratio. (i) Schematic illustrating the ratio of the major to minor axis of the fitted central peak, with increasing values indicating greater elongation. (ii) Magnitude of the change in aspect ratio from baseline for NS and EW directional representations, calculated as (Silencing+ISI)−(Baseline). Bars represent mean±SEM. Statistical comparisons were performed using paired Wilcoxon signed-rank tests; n=27 cells.

MS glutamatergic silencing induced distortions in the spatial patterns of grid maps, including smearing of the individual firing fields. To further examine these field distortions, we separated the animals’ trajectories according to their direction of travel and independently examined grid cell representations during a combination of north-south (NS) movement periods (to visualize smearing in the north-south dimension) and a combination of east-west (EW) movement periods (to visualize smearing in the east-west direction) (Figure 3E,F). Both silencing and ISI periods were combined for this analysis to ensure that we had sufficient coverage of the open field. We predicted that with disrupted velocity coding information, grid cell distortions would exhibit elongation along the corresponding axis of travel. We therefore quantified the morphology of the central peak of the spatial autocorrelation using eccentricity and aspect ratio (Figure 3G,H). The disruption of MS glutamatergic activity produced significant changes from baseline in both central-peak eccentricity and aspect ratio for NS and EW representations (eccentricity: NS, 0.13724 ± 0.0216; EW, 0.2180 ± 0.027; aspect ratio: NS, 0.5496 ± 0.3726; EW, 1.042 ± 0.3396; one-sample Wilcoxon signed-rank test against 0 displacement, *p* < 0.0001; Holm-corrected p = 2.37 10^−5; Figure 3G,H), demonstrating that silencing MS glutamatergic neurons disrupts the geometric shape of grid cell fields resulting in a distorted grid cell spatial code.

### MS glutamatergic silencing reduces strength of theta phase locking in the MEC

Given the links between septal theta generation and phase-dependent encoding mechanisms^36,47,48^, we next examined whether silencing MS glutamatergic neurons altered theta phase locking in MEC neurons relative to baseline (Figure 4A,B). To characterize heterogeneity in baseline theta coupling, we performed k-means clustering across all 44 neurons using standardized baseline MVL together with the sine and cosine of baseline preferred theta phase. The two-cluster solution showed strong separation (mean silhouette = 0.599) and identified a group of 12 neurons with higher baseline theta modulation and a group of 32 neurons with lower baseline theta modulation. Neurons in the higher-modulation cluster exhibited a reduction in MVL during silencing, from 0.253 ± 0.0129 during baseline to 0.164 ± 0.0232 during Sil and 0.175 ± 0.0368 during ISI (Figure 4C,D). MVL differed across conditions (Friedman test, *p* = 0.0183), with a significant reduction from baseline to Sil (Holm-corrected Wilcoxon signed-rank test, *p* = 0.0278). MVL did not differ between Sil and ISI (*p* = 0.970) or between baseline and ISI (*p* = 0.1040). These changes in modulation strength were not accompanied by a consistent reorganization of preferred theta phase (Figure 4E,F). Most neurons maintained their baseline phase preference during silencing (38 of 44 neurons), and the magnitude of the phase shift did not differ between the higher-and lower-modulation clusters (*p* = 0.607). Thus, MS glutamatergic silencing primarily altered the strength of neuronal coupling to the ongoing theta rhythm rather than uniformly shifting the theta phase at which MEC neurons fired. Given the role of the MS in coordinating theta oscillations and phase-dependent activity, these changes in theta-modulation strength may have important consequences for temporal coding within the MEC and downstream hippocampal networks.

**Figure 4:**
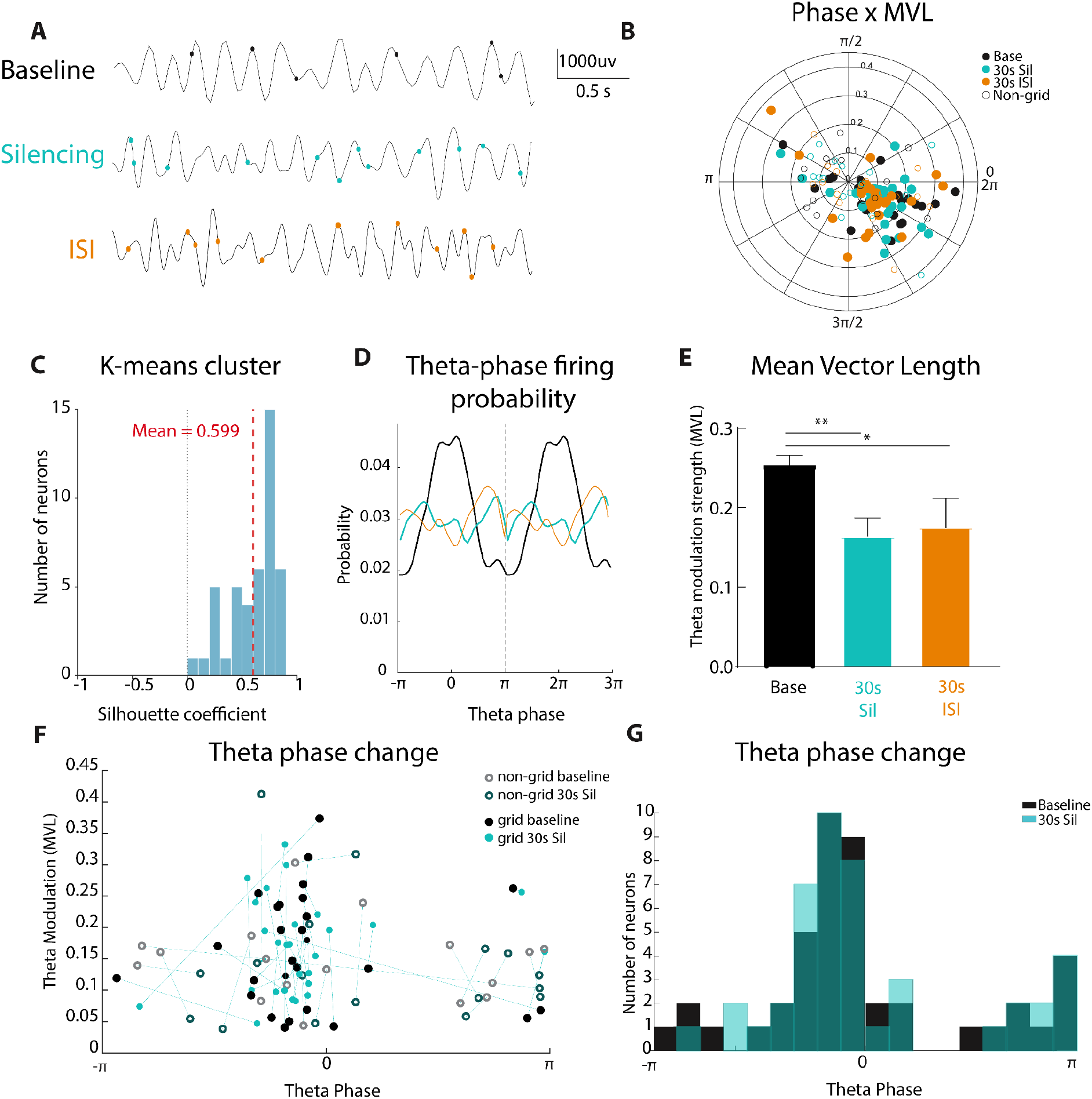
Silencing septal glutamatergic neurons reduces theta phase locking in the MEC. **(A)** Representative local field potential (LFP) traces with spike times from an individual MEC neuron during baseline (black), 30-s MS glutamatergic silencing (Sil; cyan), and the subsequent 30-s inter-stimulation interval (ISI; orange). **(B)** Polar representation of preferred theta phase and phase-locking strength for all MEC neurons (n = 44) during baseline, Sil, and ISI. Angular position indicates preferred theta phase and radial distance indicates mean vector length (MVL). Filled and open circles denote grid and non-grid cells, respectively. **(C)** K-means clustering across all 44 neurons using standardized baseline MVL together with the sine and cosine of baseline preferred theta phase. The two-cluster solution with mean silhouette indicated in red. **(D)** Theta-phase firing probability of the representative neuron shown in (A) during baseline, Sil, and ISI. Silencing reduced the concentration of firing at the preferred baseline theta phase. **(E)** Theta-modulation strength across conditions for the higher baseline theta-modulation cluster identified by k-means clustering (n = 12). **(F)** Preferred theta phase plotted against MVL for individual neurons during baseline and Sil. Lines connect measurements from the same neuron, illustrating cell-specific changes in phase preference and modulation strength. Filled and open markers indicate grid and non-grid cells, respectively. **(G)** Distribution of preferred theta phases across the population during baseline and Sil, illustrating the population-level distribution of phase preferences before and during MS glutamatergic silencing. Bars show mean ± SEM.

### Comparing experimental effects to hybrid oscillatory interference/continuous attractor network model predictions

To further understand the mechanisms underlying grid cell firing patterns, we next examined predictions from a computational grid cell model against our experimental observations. Continuous attractor network (CAN) models are currently the dominant theoretical framework for explaining grid cell spatial periodicity, whereas oscillatory interference (OI) models have received comparatively less experimental support. Rather than evaluating these models solely on their ability to generate grid-like firing, we tested whether they could reproduce the specific changes observed in our recordings following MS optogenetic manipulations. Using the experimentally observed alterations in speed coding fidelity and firing rate dynamics to guide these simulations, we simulated the effects of septal inhibition in a combined CAN/OI hybrid model and compared the resulting changes in grid cell activity to those observed in vivo.

To determine whether the experimentally observed changes in speed coding could account for the disruption of grid cell spatial periodicity, we implemented these perturbations in the hybrid oscillatory interference/continuous attractor network model of Bush and Burgess, 2014 (Figure 5). Across 100 simulated cells, reducing the slope of the speed signal significantly decreased gridness score relative to baseline (Holm-adjusted *p* = 0.0015), whereas increasing the speed slope did not significantly alter gridness (Holm-adjusted *p* = 0.1458; Figure 5C,D). We next attempted to recreate the main results of MS glutamatergic reduction by reducing speed coding fidelity, which produced the largest disruption of grid periodicity (Holm-adjusted *p* < 0.0001; Figure 5E). Notably, reduced speed coding fidelity resulted in both a loss of hexagonal periodicity and distortion of the spatial autocorrelation structure that qualitatively resembled the grid cell distortions observed during MS glutamatergic silencing and reduced model R² (Figure 3). Together, these simulations indicate that the fidelity of the speed signal can strongly influence grid cell spatial periodicity.

**Figure 5:**
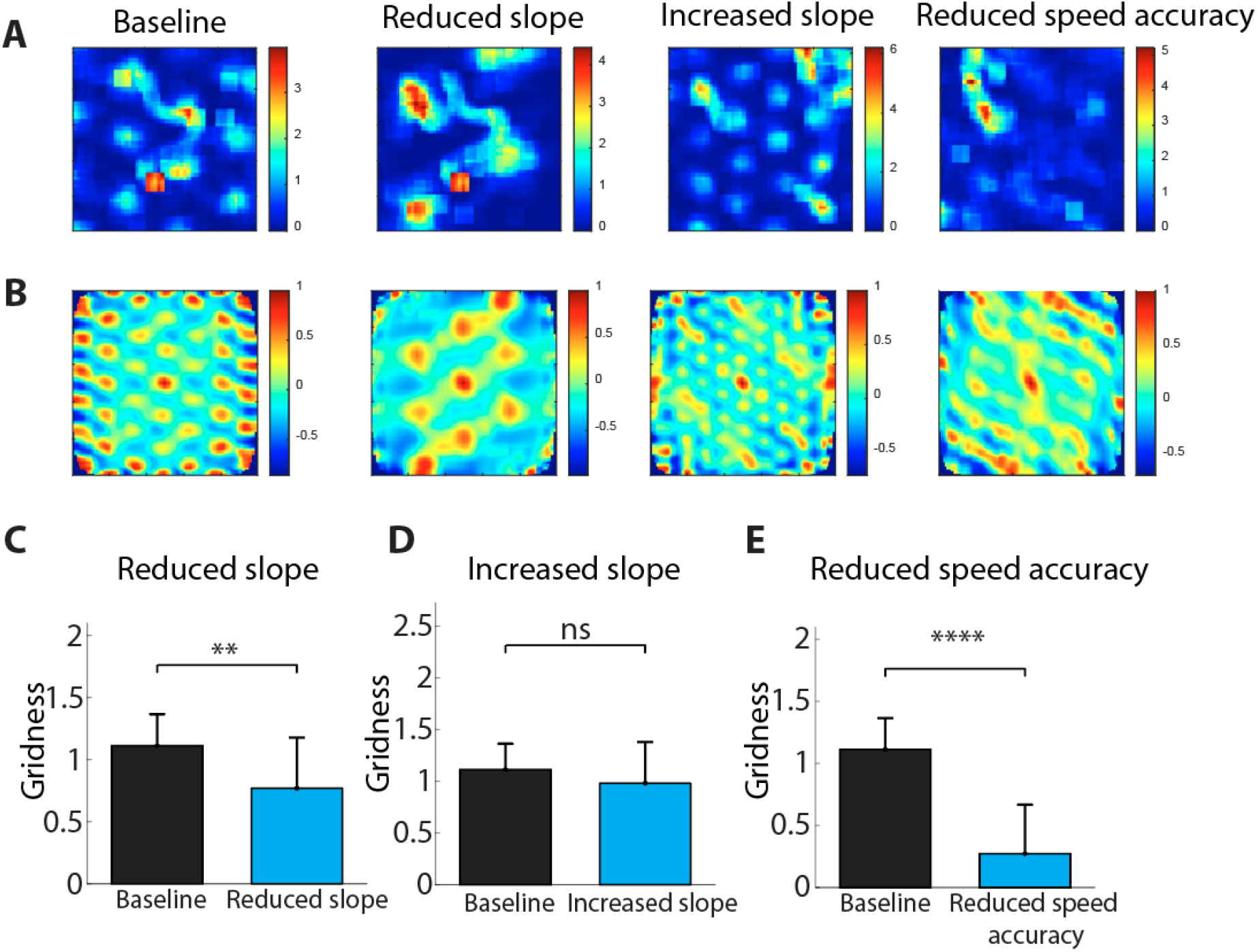
Hybrid Oscillatory Interference/Continuous attractor network model with changes to gain and speed fidelity. **(A)** Representative spatial firing-rate maps from the hybrid oscillatory interference/continuous attractor network model under baseline, reduced speed slope, increased speed slope, and reduced speed coding accuracy conditions. **(B)** Spatial autocorrelograms corresponding to the firing-rate maps shown in (A), illustrating changes in grid spatial periodicity across model conditions. **(C)** Gridness Score during baseline and reduced speed slope conditions. Reducing speed slope significantly decreased gridness relative to baseline. **(D)** Gridness Score during baseline and increased speed slope conditions. Increasing speed slope did not significantly alter gridness. **(E)** Gridness Score during baseline and reduced speed coding accuracy conditions. Reducing speed coding accuracy produced a marked decrease in gridness relative to baseline.

## Discussion

In this study, we investigated the contribution of MS glutamatergic neurons to velocity coding and grid cell spatial representations in the MEC. By combining cell-type-specific optogenetic silencing with in vivo electrophysiology, we show that silencing of MS glutamatergic neurons impairs accurate speed coding (Figure 2), theta-phase locking (Figure 4), and grid cell spatial periodicity (Figure 3). These effects occurred in the absence of marked changes in exploratory behavior or theta oscillations, suggesting that MS glutamatergic activity is not simply required for locomotion or theta generation during ongoing exploration. Instead, our findings suggest a role for MS glutamatergic neurons in maintaining movement-related signals used to organize spatial and temporal representations in the MEC.

Optogenetic activation of MS glutamatergic neurons has been associated with locomotion initiation, arousal, and behavioral state regulation, appetite suppression^49^, control of wakefulness^50^, modulation of nociception^51^, initiation of locomotion^39,44^, as well as whisking, rearing, and novel object approaches^43^. Silencing MS glutamatergic neurons reduced exploration during anxiogenic conditions^43^, however, in our hands, silencing this population does not appear to cause marked changes in exploration when animals were in a familiar environment for free foraging. This dissociation suggests that an important function of MS glutamatergic neurons is to convey movement-related information to downstream circuits, not necessarily for movement initiation itself.

Grid cell spatial periodicity depends on the integration of multiple sources of information, including visual cues^52^, directional signals from the anterior thalamic system^52,53^, excitatory drive from the hippocampal formation^54^, and rhythmic and velocity-related inputs from the MS^17,27^. Consistent with a disruption of movement-dependent signals, MS glutamatergic silencing reduced grid cell spatial periodicity with distortions in the geometry of individual grid fields. Direction-conditioned rate maps and spatial autocorrelations revealed elongation of the grid representation, suggesting that the perturbation was expressed within the spatial structure of the grid code rather than solely as a reduction in gridness. One possibility is that they reflect the accumulation of error within the path-integration system. Grid representations accumulate spatial error as a function of time and distance travelled, whereas encounters with environmental boundaries provide external spatial information capable of correcting this accumulated error and re-anchoring the grid representation^55^. Within this framework, reducing the accuracy of velocity information during MS glutamatergic silencing could increase the discrepancy between the animal’s actual movement and the displacement represented by the grid network. The resulting error would be expected to accumulate during movement until corrected by re-anchoring, potentially contributing to the elongation and smearing distortions and instability of grid cells observed here in Figure 3.

In addition to disrupting speed and spatial coding, MS glutamatergic silencing altered theta phase locking in the MEC. This suggests that MS glutamatergic neurons contribute to the temporal organization of MEC activity within the theta cycle. Theta oscillations are predicted to provide a temporal scaffold to organise the activity of spatially tuned neurons within the hippocampal network, where different phases of the theta cycle are proposed to support distinct computational roles. Late phases of theta are associated with prospective spatial representations via phase precession, whereas early phases have been hypothesized to support retrospective spatial representations and the encoding of new associations. Consistent with this temporal organization, long-term potentiation and long-term depression in the rodent hippocampus depend on theta phase, suggesting that the timing of inputs relative to the theta cycle influences synaptic plasticity and memory encoding^56–59^. The observation that MS glutamatergic silencing alters the mean vector length while largely preserving preferred theta phase suggests that MS glutamatergic neurons may regulate the strength with which MEC neurons are temporally coupled to the ongoing theta rhythm. Furthermore, the convergence of effects on speed coding and theta phase locking raises the possibility that MS glutamatergic input may be particularly important for computations that require movement information to be organized within the theta cycle. Theta sequences and prospective sweeps^60^ depend on the sequential activation of spatial representations at compressed timescales, and their spatial extent must ultimately be related to movement through the environment. A pathway capable of influencing both the fidelity of speed-related activity and the strength of theta phase locking is therefore well positioned to coordinate these spatial and temporal dimensions of MEC activity. MS glutamatergic neurons may therefore influence temporal processes relevant to theta phase precession, prospective sweeps, and memory.

Together, our results identify MS glutamatergic neurons as an important component of the circuit mechanisms that couple self-motion to spatial representations in the MEC. Rather than simply controlling locomotion or theta activity, MS glutamatergic neurons support the fidelity of speed-related firing, the coupling of MEC neurons to theta, and the stability of grid cell spatial representations. These findings refine our understanding of the signals required to maintain an internally generated cognitive map and suggest that stable spatial representations depend on the coordinated contribution of multiple, functionally distinct septal inputs. Disruption to this system may provide insight into how septo-entorhinal circuit dysfunction contributes to impaired spatial navigation in neurological disease.

## Methods

### Subjects

All procedures were performed according to protocols and guidelines approved by the Institutional Animal Care and Use Committee at Boston University and the McGill University Animal Care Committee and the Canadian Council on Animal Care. VGLUT2^Cre^ knock-in homozygote mice (The Jackson Laboratory, stock # 016962) were housed in a 12:12 hour light/dark cycle with food and water ad libitum. Mice were injected between 8-12 weeks of age and implanted 2-3 weeks following injection. Mice were recorded for between 4-6 weeks. Both male and female mice were implanted however, only male mice are included in the current dataset due to coverage issues in female mice. The housing room conditions of the mice were maintained at 20–22 degrees Celsius and 21–30% humidity. Specificity of this mouse line for Cre recombinase has previously been confirmed in this mouse line^40^.

### Surgeries

For all surgeries, mice were anesthetized via oxygen and 5% isoflurane in an inhalation box and then transferred to the stereotaxic frame (David Kopf Instruments), where anesthesia was maintained via inhalation of oxygen and 1-2.5% isoflurane for the duration of the surgery. The animals’ body temperature was maintained via heating pad and eyes were protected with hydrogel (Optixcare). The analgesic carprofenThe anesthetic Carprofen (10 ml kg^-1^) and saline (0.5 ml) were administered subcutaneously before each surgery.

To trigger the expression of the opsins, VGLUT2^Cre^ mice were stereotaxically injected in the MS with a Cre-dependent adeno-associated viral vectors. Silencing experiments were performed with ArchT-GFP (AAVdj-EF1α-Flex-ArchT-GFP from University of North Carolina Virus Core) and virus control experiments were performed with GFP (AAV2-syn-GFP from). Viruses were delivered directly into the medial septum at AP 0.86 mm from bregma, ML 0.0 mm, DV 4.5-4.7mm. All injections were administered via glass pipettes connected to a Nanoject II (Drummond Scientific) injector at a flow rate of 23 nl/sec. Following a 2-4 week incubation period, mice underwent a second surgery to implant an optic fibre and microdrive. For each mouse, two stainless anchor screws (B000FN0J58, Antrin Online) were placed in front of the inferior cerebral vein, and were secured to the skull with dental cement (Patterson Dental, Inc). A ground wire was positioned above the contralateral cerebellum. For optogenetic modulation of the medial septum an optic fiber (CF230, Thorlabs) connected to a ferrule (Precision Fiber Products or Thorlabs) was implanted at a 5° angle to target just above the medial septum (AP 0.86, ML 0.2, DV 3.83 mm) and secured to the skull with dental cement. Electrophysiological experiments were performed with either custom made microdrives. The microdrive used consisted of four independently movable tetrodes (Axona, Inc) and was implanted above of the MEC at the following stereotaxic coordinates: 3.4 mm lateral to the midline, 0.25-0.40 mm anterior to the transverse sinus at an 8-10°angle. Tetrodes were gold-plated using the NanoZ (Neuralynx) to lower impedances to 200-250 kΩ at 1 kHz and dipped in mineral oil prior to surgery. The microdrive was secured in place with metabond and dental cement (Patterson Dental). Once the cement was dry, tetrodes were lowered 400um below the surface of the brain. Mice body temperature was maintained using a heating pad throughout the surgery until fully recovered.

### Data acquisition

Following surgery, animals had 1 week of recovery. Once recovered, mice were placed on water restriction and maintained at 85% of their ad libidum weight for the duration of experiments. Animals were connected to custom-built headstage preamplifier tethers (Neuralynx) and an optic fiber patch cord (Thorlabs). Light delivery was achieved through an optic fiber patch cord and sleeve coupled to 520nm wavelength Doric Laser with a maximum power output: 20mW. Neural recordings were amplified and band-pass filtered between 0.6 kHz and 6 kHz using a Digital Lynx SX recording system (Neuralynx) and stored with Cheetah Software (Neuralynx). Animals were recorded in an open field (75 x 75 cm square box). Tetrodes were advanced at 25 µm increments to sample neurons. Spike waveform thresholds were adjusted before commencing each recording and ranged between 25-140 µV depending on unit activity. Waveforms that crossed the threshold were digitized at 32 kHz and recorded across all four channels of the given tetrode. Local field potentials were recorded across all tetrodes.

### Optogenetic Experiments

During recording, the environment was dimly lit and white-noise was played to mask uncontrolled ambient sounds. Following each recording the environment was cleaned with disinfectant (Prevail). Once grid cells had been identified during a baseline recording light delivery was achieved through an optic fiber patch cord and sleeve coupled to the laser and light intensity at the tip of the optic fiber was estimated for between 15-20 mW. Stimulation protocols consisted of 30s laser on periods followed by 30s laser off periods. The present dataset includes recordings from 19 recording sessions from four mice which were split into ArchT vs Laser control groups (13 vs 6 session respectively).

### Histology

Immunocytochemistry was used to assess the virus distribution across the septum and localize tetrode recording sites in the MEC. Experimental animals were euthanized following experiments. Mice were anesthetized and intracardially perfused with 4% paraformaldehyde in PBS. Brains post-fixed by immersion in the same fixative for 2 weeks with the tetrodes in place following perfusion and then dissected from the skull. Free-floating coronal sections of the entire medial septum and sagittal sections of the entire entorhinal cortex were cut using a vibratome (40 μm) or on the cryostat (25 μm). Free-floating sections across the septum were incubated in GFP rabbit antiserum (1:1000, Invitrogen), and were detected with anti-rabbit coupled to AlexaFluor-488 (1:1000, Invitrogen). Sections of the entorhinal cortex were stained for DAPI to locate tetrode tips. The slices were mounted with Fluoromount-G (Southern Biotechnology) and analyzed with an AxioObserver.Z1 microscope (Carl Zeiss).

### Data processing

Cluster cutting. Single-units were isolated ‘offline’ manually using graphical cluster cutting software (Offline Sorter, Plexon Inc.) individually for each recording session. Neurons were separated based on the peak amplitude and principal component measures of spike waveforms. Stability of units was confirmed by tracking waveform profiles across the four leads of the tetrode and cluster position across recording sessions, comparing to the baseline session. Position estimation. To estimate the position of the animal, we measured the centroid of a group of red and green diodes positioned on the recording head stage. Head direction was calculated as the angle between the red and green diodes.

## Data analysis

To analyze LFP power, the power spectrum for local field potentials was obtained using multitaper method included in the Chronux toolbox^61^ (mtspectrumc with NW = 3 and K = 5. Theta power was calculated by taking the area within 1Hz of the maximum power in the theta range (6-10Hz). Baseline theta power was obtained by taking the average of theta power during laser-off trials, and mean reduction in theta power for each session was obtained by dividing each laser-on trial with baseline theta power and averaging across laser-on trials. For each channel, the theta-delta ratio was obtained by taking the ratio between mean theta power (6-12Hz) and mean delta power (2-4Hz) during laser-off segments. The channel with the highest theta-delta ratio was used in the LFP analysis.

Speed–theta relationship: Running speed was calculated from successive position samples and converted to cm/s and smoothed with a Kalman filter. Continuous theta power was obtained by band-pass filtering the LFP between 4–12 Hz and calculating the squared magnitude of the Hilbert transform. Theta power was interpolated to the position-sampling timestamps and averaged within 1 cm/s speed bins. The relationship between running speed and theta power was quantified over a common speed range for laser ON and OFF periods, extending from 2.5 cm/s to the speed at which the smoothed laser-OFF theta-power relationship reached its maximum. Linear regression was used to quantify the slope and coefficient of determination (*R*²) of the speed–theta relationship.

### Measurement of single unit properties

#### Speed score

The overall correlation between the running speed of the animal and the firing rate of the individual MEC cells following the same procedures as Kropff et al. (2015). Briefly, the spike times for each cell were first binned (30ms) and then convolved with a gaussian window (300ms; 30ms s.d.). The resulting continuous firing rate was then interpolated to match the running speed vector and a speed score using the Pearson correlation was computed. A shuffle surrogate set was generated by randomly circularly shifting the speed vector relative to the firing rate with a minimum offset of 30s. A cell was considered to be speed modulated if it exceeded the 99th percentile of the shuffle distribution.

#### Speed profile

To determine if the speed tuning followed a linear or saturating profile (Hinman et al. 2016) the firing rate data from each frame with a velocity greater than 2.5cm/s was binned (1cm/s) up to the 90th percentile of the running speed for that session. The binned rate (FR) was then fit to two functions: linear (FR = b • velo + a) or a saturating exponential (FR = d - a • e^-b•velo^) using the ‘fit’ function in MATLAB. The speed profile for the cell was determined based on the model with the higher R^2^.

#### Spatial coding analysis

Spatial rate maps were constructed for a given unit by taking the number of spikes in 3.6 cm by 3.6 cm spatial bins and dividing by the amount of time spent in that bin. Rate maps were smoothed using a 5×5 bin 2D-Gaussian kernel with a one-bin standard deviation. The spatial periodicity of grid cells was quantified with a ‘‘gridness” score and computed from the spatial autocorrelation of the smoothed rate maps^27^. In brief, to calculate the gridness score, ‘‘gridness 3” from Brandon et al. (2011), the center peak was removed from the autocorrelation and if present, the six surrounding peaks were found and cut out to make a donut. If the grid shape was elliptical, it was distorted to create a circle. The correlation was calculated for each 3-degree rotation of the donut to itself. The gridness score is the difference between the correlation at the minimum peak of 60 or 120 degrees and the maximum trough at 30, 90, or 150 degrees. In line with previous work^44^, grid cells were defined as cells with a grid score >=0.19 and at least 300 spikes in the baseline condition. The spatial information of a cell was calculated using the same rate map as above, with the equation for I in bits/spike where p_i_ is the probability of occupancy in pixel i, F_i_ is the firing rate for pixel I, and F is the mean firing rate.

To quantify changes in the geometry of grid-cell spatial representations, spatial autocorrelations were analyzed separately for north–south (NS) and east–west (EW) directional representations. Cells were included only when measurements were available for both directions during baseline and MS glutamatergic silencing. The central autocorrelation peak was characterized using two morphological measures: eccentricity and aspect ratio. For each cell and direction, the magnitude of the stimulation-induced change was calculated as the absolute difference between stimulation and baseline, ∣Stim−Baseline∣|\mathrm(Stim)-\mathrm(Baseline)|, such that zero represented no change and increasing values represented greater deviation from the baseline spatial autocorrelation geometry.

#### Theta phase modulation

Theta phase was extracted from the LFP and interpolated to the timestamp of each spike. Spike phases were binned from −π to π in π/16-radian bins, and theta phase-locking strength was quantified for each neuron using the mean resultant vector length (MRL). The circular mean was additionally calculated to determine each neuron’s preferred theta phase. These measures were calculated independently during baseline, MS glutamatergic silencing, and ISI periods. For visualization, spike-phase distributions were converted to probability distributions and smoothed with a Gaussian kernel.

#### Hybrid Model

The hybrid model was modified from Bush & Burgess, 2014 hybrid model, which was obtained from https://modeldb.science/218085?tab=2. We then altered the script to independently manipulate the gain and fidelity of the velocity signal used to update the continuous attractor network. To alter the slope of the speed-dependent input while preserving the underlying trajectory, we scaled the velocity gain parameter, α\alpha, relative to baseline: reduced-slope simulations used 60% of the baseline gain (α×0.6\alpha \times 0.6), whereas increased-slope simulations used 140% of baseline (α×1.4\alpha \times 1.4). To reduce speed-coding accuracy without changing gain, we instead corrupted the velocity estimate supplied to the network. Cumulative Gaussian noise was added to the animal’s true x-and-y-position trajectories, with noise amplitude determined by 1−speedAccuracy1-\mathrm(speedAccuracy); thus, baseline simulations used an uncorrupted trajectory (speedAccuracy=1.0\mathrm(speedAccuracy)=1.0), whereas reduced-accuracy simulations used speedAccuracy=0.6\mathrm(speedAccuracy)=0.6. Effective velocity was then calculated from the corrupted trajectory and used as the movement-related input to the network, while the true trajectory was retained for subsequent spatial analyses. This approach allowed changes in velocity gain and velocity-signal fidelity to be examined independently. The resulting firing rate–speed relationship was quantified after simulation, rather than fixing the model to a predetermined R² value.

## Statistics

Normality of distributions was not assumed, so comparisons were made using non-parametric statistics. All statistical tests are noted where the corresponding results are reported throughout the main text and supplement. All tests were 2-tailed unless otherwise noted. Nonparametric tests were used as neural data violates the assumptions of parametric tests.

All statistical evaluations were performed under MATLAB and Prism. Non-parametric Wilcoxon signed rank tests, Wilcoxon rank sum tests, Kolmogorov-Smirnov tests, and Kruskal-Wallis or Friedman test one-way ANOVAs were used throughout the paper. P-values reported from all post hoc tests were with Dunn’s multiple comparison test. Comparisons of single distributions for a difference from zero were performed using Wilcoxon sign-rank tests. For comparisons of two distributions Wilcoxon sign-rank or Wilcoxon rank-sum tests were used.

## Supplemental Figures

**Supplementary Figure 1.**
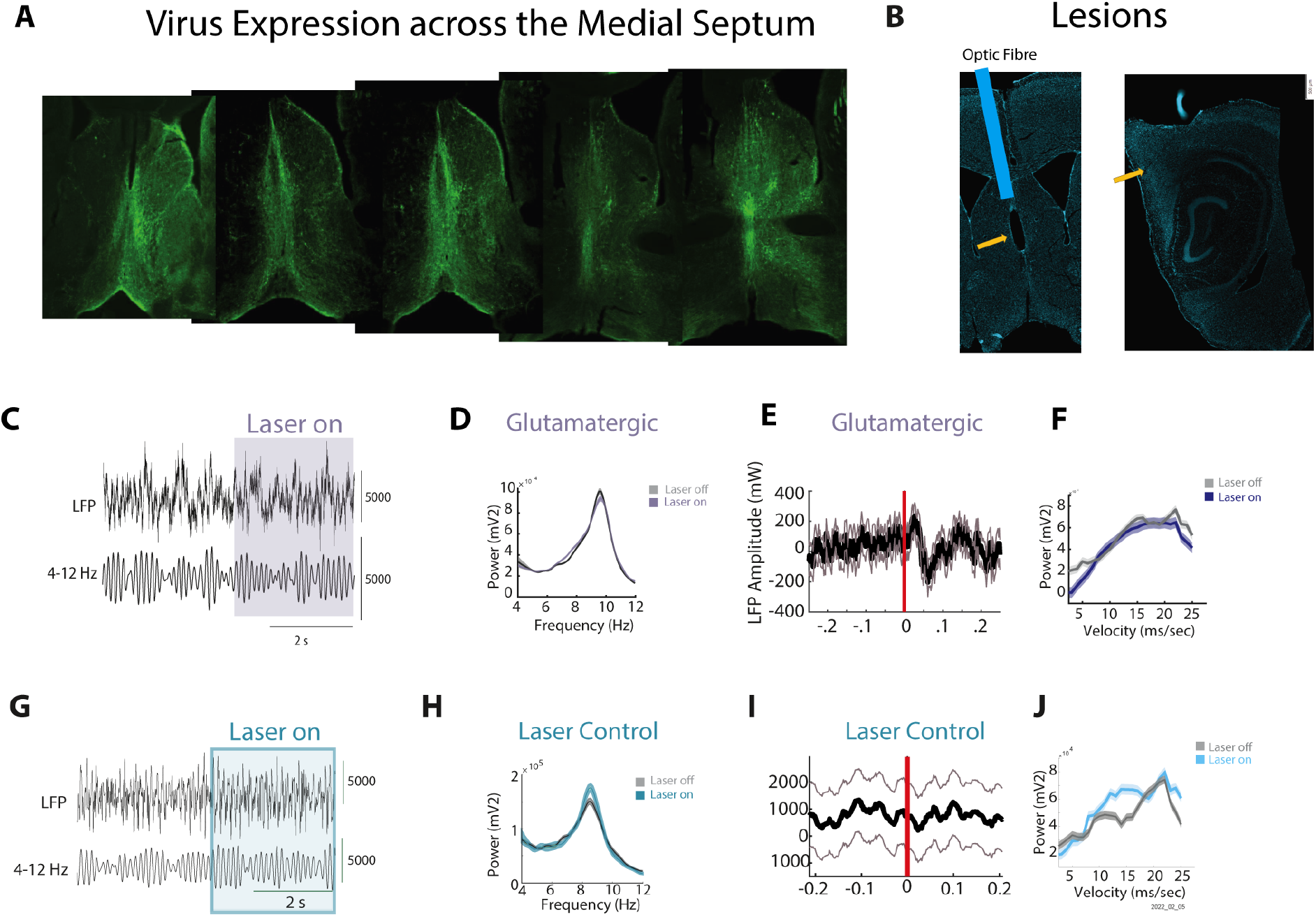
Histological verification and laser control analyses of LFP recordings in the MEC. (**A**) Representative coronal sections spanning the medial septum illustrate viral expression across the anterior–posterior extent of the targeted region. (**B**) Representative histological verification of the optic-fibre tract in the medial septum (left; blue overlay) and recording-site lesion in the hippocampal formation (right); yellow arrows indicate lesion locations. (**C**) Representative raw local field potential (LFP; top) and theta-band-filtered signal (4–12 Hz; bottom) recorded during inhibition of MS glutamatergic neurons. Shading indicates the laser-on period. (**D**) Mean LFP power spectrum during laser-off (grey) and laser-on (purple) periods in the MS glutamatergic silencing group. (**E**) LFP amplitude aligned to laser onset (red vertical line) in the glutamatergic group. The black trace shows the mean and the flanking traces indicate variability across observations. (**F**) Relationship between running velocity and LFP power during laser-off (grey) and laser-on (purple) periods in the glutamatergic group. (**G**) Mean power spectrum during laser-off and laser-on periods for laser control group (**H**) laser-onset-aligned LFP amplitude (**I**) Relationship between running velocity and LFP power during laser-off (grey) and laser-on (blue) periods in the control group

**Supplementary Figure 2.**
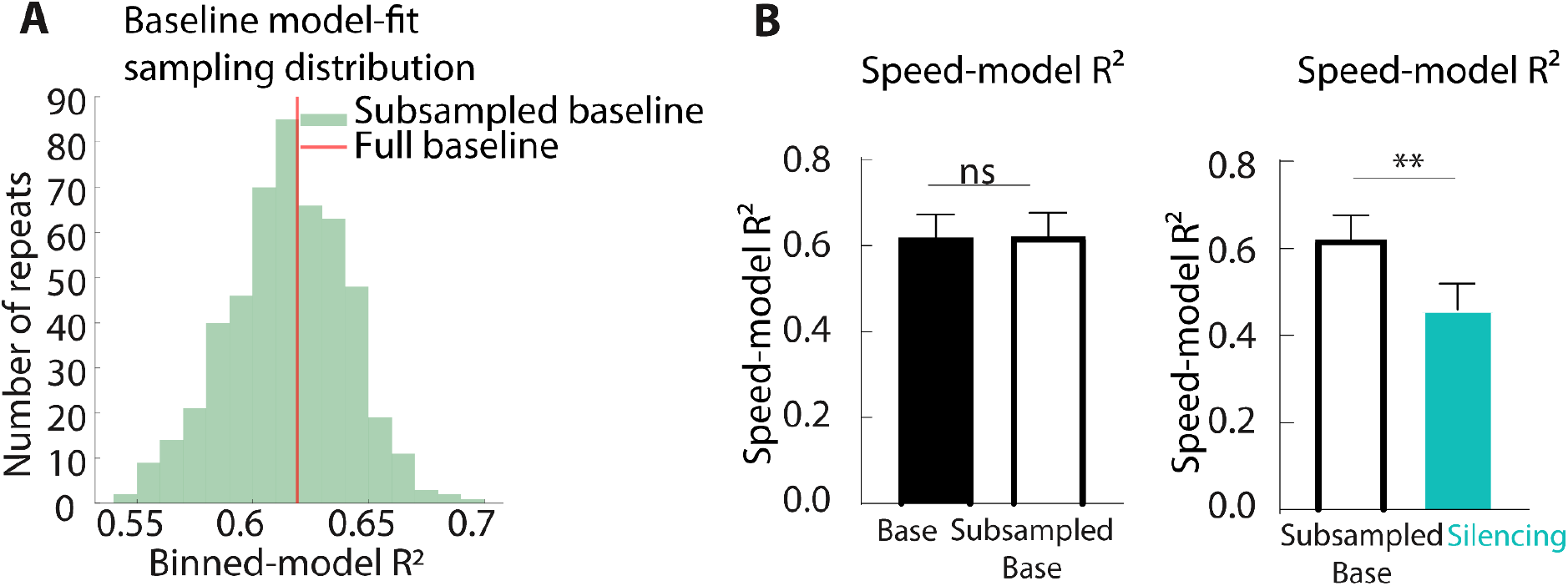
Baseline subsampled to the same number of 30s epochs as contained in corresponding Silencing period. **(A)** Sampling distribution of the mean winning binned speed-model R² obtained from 500 randomly generated Baseline subsamples. For each recording session, each subsample contained the same number of non-overlapping 30-s epochs as the number of 30-s Laser-on epochs from between 2.5-45cm/s. The red vertical line indicates the mean winning binned-model R² calculated from the full Baseline period. **(B)** Speed-model R² for the full Baseline and the median subsampled Baseline value for each cell (left), and for the subsampled Baseline and Silencing periods (right). Subsampling did not significantly alter Baseline model fit (*p =* 0.4980), whereas speed-model R² was significantly lower during Silencing than during the matched subsampled Baseline (*p* < 0.01). Bars in B show mean ± SEM. ns, not significant; \*\**p* < 0.01.

**Supplementary Figure 3.**
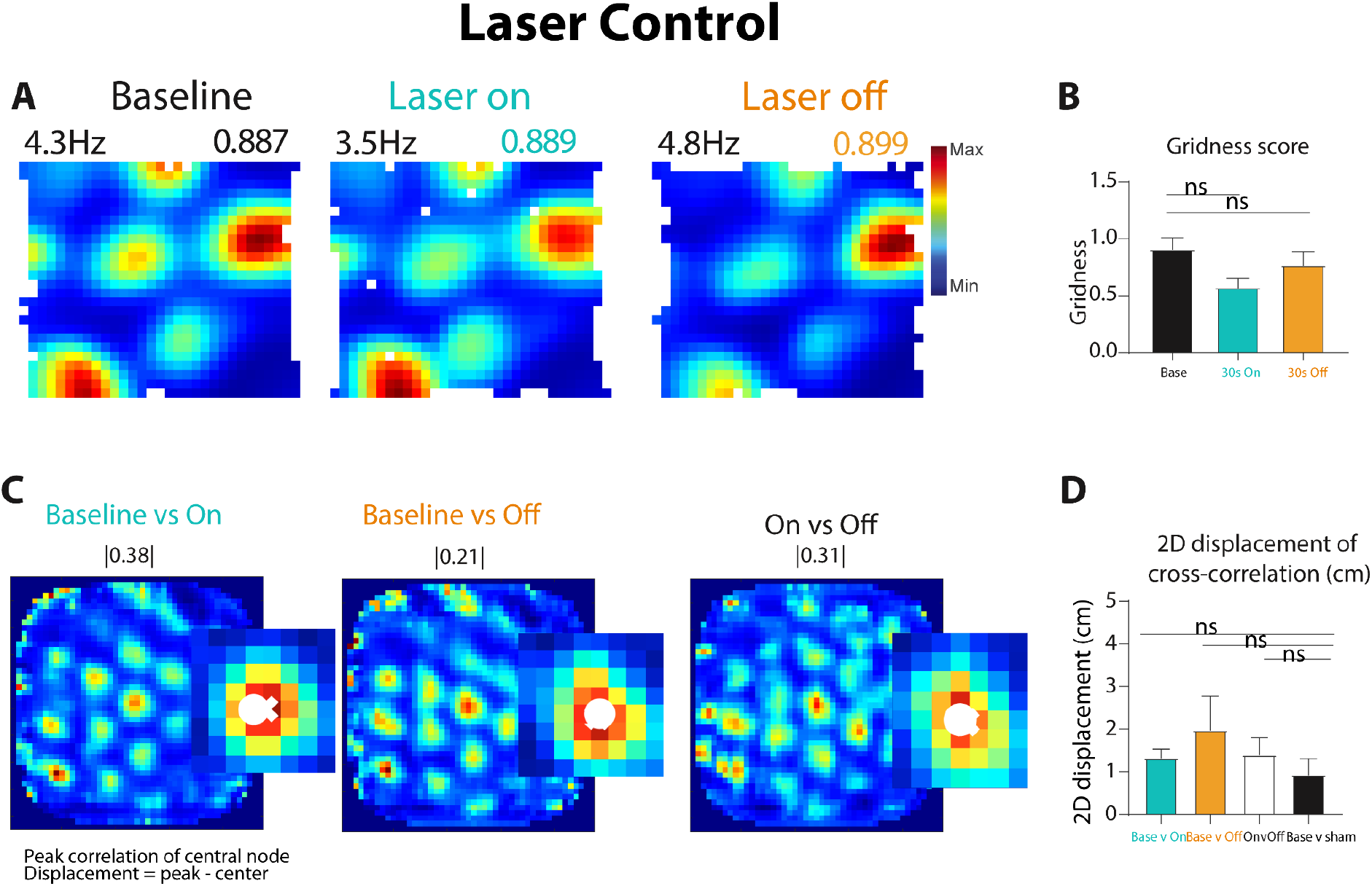
Grid cell laser control experiments across baseline, laser on and laser off conditions. **(B)** Representative spatial firing-rate maps from a grid cell recorded in a Laser Control animal during Baseline, 30-s laser-on periods, and the 30-s laser-off periods. Peak firing rate (Hz; upper left) and gridness score (upper right) are shown for each condition. **(C)** Mean gridness scores during Baseline, Laser On, and Laser Off. Gridness scores did reduce slightly with the laser on however it did not differ between Laser On and Baseline (Dunn’s multiple-comparisons test, adjusted *p* = 0.7158) or between Baseline and ISI (*p* > 0.9999). **(D)** Spatial cross-correlations for Baseline versus Laser On, Baseline versus Laser Off, and Laser On versus Laser Off. Insets show the central cross-correlation peak, with the white circle indicating its location relative to the centre. Values above the maps indicate the magnitude of the peak displacement from the centre. **(E)** Mean two-dimensional displacement of the central cross-correlation peak for Baseline versus Laser On, Baseline versus Laser Off, Laser On versus Laser Off, and Baseline versus sham comparisons. No pairwise comparison was significant (Dunn’s multiple-comparisons test: Baseline–Laser On versus Baseline–Laser Off, adjusted *p* > 0.9999; Baseline– Laser On versus Laser On–Laser Off, *p* > 0.9999; Baseline–Laser On versus Baseline–sham, *p* = 0.5861; Baseline–ISI versus Laser On–Laser Off, *p* > 0.9999; Baseline–Laser Off versus Baseline–sham, *p* = 0.2306; Laser On–Laser Off versus Baseline–sham, *p* = 0.5861). Error bars indicate SEM; ns, not significant.

